# Characterisation and correction of polarisation effects in fluorescently labelled fibres

**DOI:** 10.1101/2023.11.10.566047

**Authors:** Nandini Aggarwal, Richard Marsh, Stefania Marcotti, Tanya J Shaw, Brian Stramer, Susan Cox, Siân Culley

## Abstract

Many biological structures take the form of fibres and filaments, and quantitative analysis of fibre organisation is important for understanding their functions in both normal physiological conditions and disease. In order to visualise these structures, fibres can be fluorescently labelled and imaged, with specialised image analysis methods available for quantifying the degree and strength of fibre alignment. Here we show that fluorescently labelled fibres can display polarised emission, with the strength of this effect varying depending on structure and fluorophore identity. This can bias automated analysis of fibre alignment and mask the true underlying structural organisation. We present a method for quantifying and correcting these polarisation effects without requiring polarisation resolved microscopy, and demonstrate its efficacy when applied to images of fluorescently labelled collagen gels, allowing for more reliable characterisation of fibre microarchitecture.

## Introduction

Many biologically important proteins form linear filaments, fibrils and fibres. Studying such structures *in vitro, ex vivo* and *in vivo* typically relies on visualisation using either intrinsic fluorescence properties of these proteins or labelling them with exogenous fluorophores. Many of these proteins play structural roles, and as a result analysis of their collective organisation can provide insight into cell and tissue mechanics.

Fluorescence labelling of biological structures, fibrous or otherwise, requires three components: the target biological molecule to be labelled, a fluorescent molecule (fluorophore), and a linker molecule connecting the two. For labelling with fluorescent proteins, the linker molecule is either a short sequence of residues between the target protein and the fluorescent protein sequence, or is absent with the fluorescent protein sequence fused directly to the target sequence. When labelling with small organic dye molecules, linkers can be small covalent linking molecules (e.g. NHS esters and other click chemistry approaches (1)), target-specific molecules (e.g. phalloidin for actin labelling) and antibodies/nanobodies for immunofluorescence. The combination of target molecule, linker, and fluorophore can lead to differences in properties of fluorescence emission when illuminated with polarised light.

When light interacts with a fluorophore, the probability of excitation is dependent on the angle between the electric field vector **E** (polarisation) of the incident illumination and the absorption transition dipole moment *μ*_ab_ of the fluorophore. For linearly polarised single photon excitation this probability follows a cos^2^ θ distribution, where θ is the angle between **E** and *μ*_ab_; molecules with *μ*_ab_ oriented parallel to **E** will be efficiently excited, whereas molecules with *μ*_ab_ oriented perpendicular to **E** will not be excited. The polarisation of fluorescence emitted by a molecule is similarly polarised parallel to the emission transition dipole moment *μ*_em_ (which is usually close to parallel to *μ*_ab_) (2). In conventional confocal and widefield fluorescence microscopy applications, polarisation effects are typically neglected. This is due to assumptions that fluorophores are, if fixed, randomly oriented or undergoing free rotational diffusion which changes their orientation on a (typically) sub-nanosecond timescale (Figure 1a). Specifically, it is assumed that fluorophore reorientation is fast compared to the fluorescence lifetime, a condition easily met for unbound fluorophores in an aqueous environment. This is in contrast to the situation where fluorophore alignment or rotation is in some way constrained by the structure, linker and/or fluorophore (Figure 1b).

**Fig. 1.**
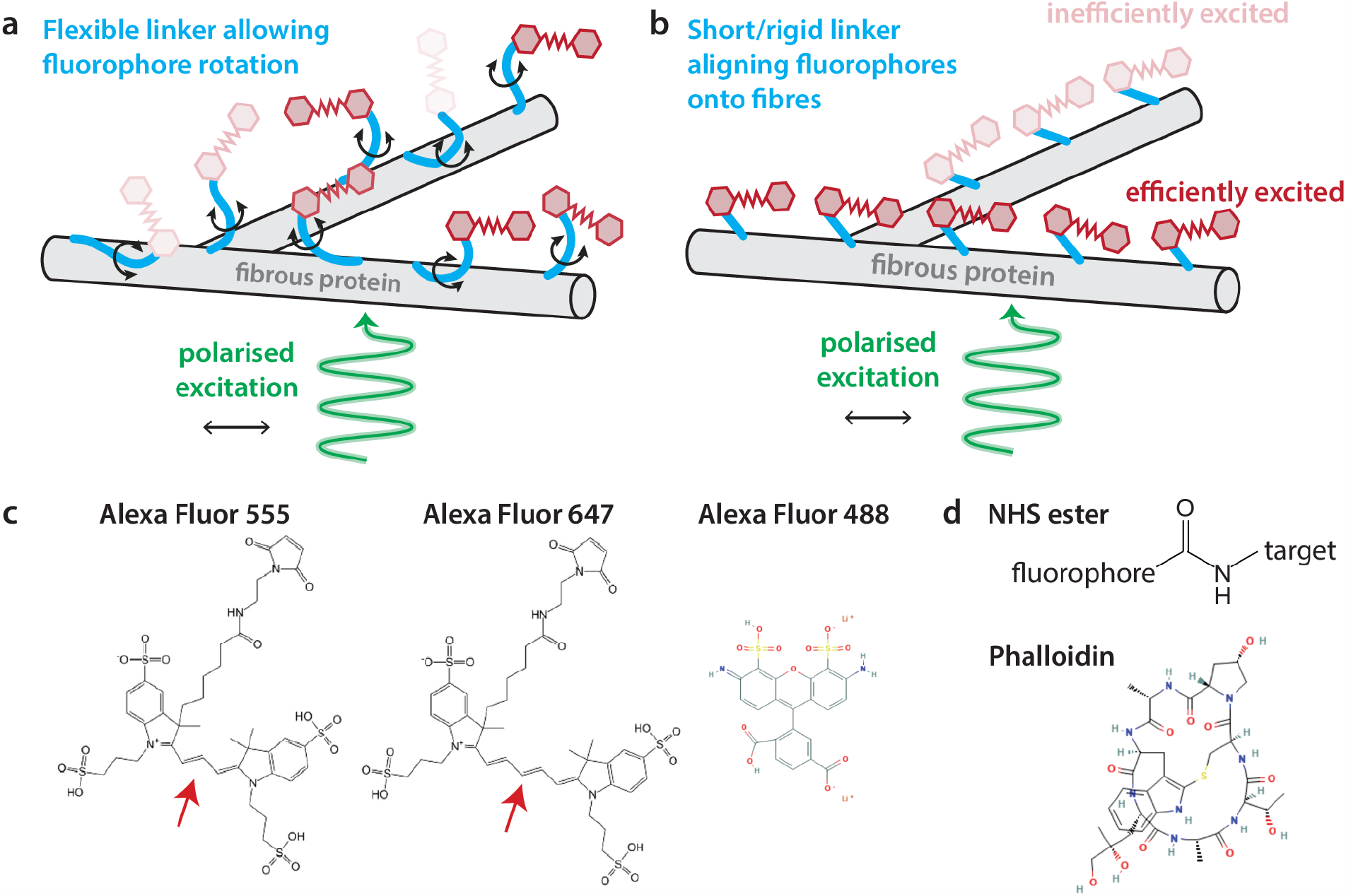
Contributing mechanisms to unpolarised and polarised emission from fluorescently labelled fibres. **a** An example of assumed label behaviour when fibrous structures are labelled with fluorescent molecules (red shades) via a linker molecule (light blue). Here, rotational diffusion of the labelling complex is unrestricted and faster than the fluorescence lifetime of the fluorophore, preventing long-term alignment of excited fluorophores along the axis of the fibres. As a result, even if illuminated by polarised light (green, arrow indicates polarisation direction), the random orientations of the fluorophores relative to the structure when emitting would result in no net polarisation of the fluorescence (darkness of red indicates probability of fluorophore being excited). **b** A mechanism by which the system presented in **a** could generate polarised emission. A combination of fluorophore structure (temporary or permanent direct binding to the fibre) and/or small, rigid linker molecule (restricted orientational freedom) could lead to alignment of fluorophores along the axes of fibres. Excitation with polarised illumination here would lead to efficient excitation of fluorophores on fibres parallel to the polarisation direction (dark red) and as a result more emitted fluorescence when compared to fluorophores at a larger angle from the polarisation direction (pale red). **c** Structures of fluorophores Alexa Fluor 555, Alexa Fluor 647 (both taken from (5)) and Alexa Fluor 488 (6). Arrow indicates difference in structure between Alexa Fluor 555 and Alexa Fluor 647. **d** Structure of NHS ester linkage and phalloidin (7) used as linkers.

The relationship between fluorescence lifetime *(τ*_*f*_) and rotational diffusion time (*τ*_rot_) in relation to the anisotropy of emitted fluorescence is described by the Perrin equation (2)

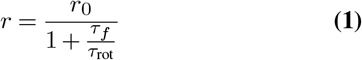

where *r* is the measured fluorescence anisotropy and *r*_0_ is the fundamental fluorescence anisotropy without any rotation. The assumption that the rotational diffusion time is much faster than the fluorescence lifetime (*τ*_rot_ *≪ τ*_*f*_) ensures that, regardless of any intrinsic polarisation in the fluorophore, the measured anisotropy is 0 (i.e. there are no observed polarisation effects). While *τ* _*f*_ depends predominantly on the fluorophore identity, *τ* _*rot*_ depends on both fluorophore identity and linker. For example, large, bulky fluorophores such as GFP will have longer *τ* _rot_ compared to small organic dye molecules, and shorter, stiffer linkers may yield longer *τ* _rot_ compared to longer, floppier linkers. This can be exploited in experimental design, for example when GFP is inserted into rigid domains of integrins for probing their orientation (3, 4). Here, presence/absence of a short linker and truncation of the GFP sequence alter the constraint of GFP rotation and as a result the measured anisotropy.

Polarisation effects are often considered for specific applications such as Förster Resonance Energy Transfer (FRET) microscopy, where orientation restriction between donor and acceptor dipoles can bias measurements of FRET efficiency (8). Fluorescence polarisation in itself can also be used as a powerful low-invasive and label-free imaging tool for analysing the order of cellular and molecular level tissue architecture of both hard (9) and soft (10) tissue. However, for conventional intensity-based fluorescence microscopy studies, polarisation effects are rarely considered. This can pose problems as, even if an experiment has not been specifically designed to use or observe polarisation, polarisation effects will frequently be present within fluorescence microscopy data. The huge number of combinations of target structure, labelling method, fluorophore and imaging system in biological microscopy also make it difficult to predict what polarisation effects may be present.

Here, we are concerned with imaging fluorescently labelled fibre structures and analysing their organisation. There are two main approaches to quantifying the organisation of fibrillar structures in microscopy images. One approach is to segment individual fibres and analyse properties of these segmentations, while the other is to examine features of images such as local intensities and gradients. FibrilJ (11) and CT-FIRE (12) are commonly-used examples of the first approach; while such approaches are successful for electron microscopy imaging modalities, threshold-based methods often perform poorly on fluorescence microscopy data where both structure and background have more variable intensities, and are often computationally intensive. Therefore, most fibre analysis software used for analysing fluorescence microscopy data is image-based. Methods such as FibrilTool (13) and OrientationJ (14–16) use image processing with nematic tensors and structure tensors respectively to extract information about local order and orientation, based on calculation of local intensity gradients. Alternatively, images can be analysed in frequency space by analysis of the Fourier spectrum of the image. This can be done on the whole image (CytoSpectre (17)) or in a patchwise manner that allows for quantification of alignment at different length scales (Alignment by Fourier Transform, AFT (18)). Although these methods cannot provide information on individual fibres, they quantify bulk behaviour of fibres within a region of interest. These segmentation and bulk analysis approaches are combined in TWOMBLI (19), allowing for analysis of both individual fibre and global patterns.

One example of a fibre-forming protein whose organisation relates to function is collagen. Collagens are the most abundant proteins in humans, constituting approximately one third of total protein mass (20). To date, 28 different types of collagens (I-XXVIII) have been identified (21), of which type I has the largest distribution (22). Collagen type I is a fibrillar collagen that is commonly found in the skin, tendons and bones. The dominant macromolecule present within the extracellular matrix (ECM), collagen type I is composed of three long polypeptide chains wound together tightly to form a stable triple helix structure. These triple helix structures are then assembled into fibrils and fibres that act to provide support, mechanical strength and structure to surrounding cells and tissues. Due to its important role, studies have shown that alterations to collagen fibrillar alignment result in matrix changes that can significantly affect tissue integrity and contribute to the genesis and progression of numerous pathological conditions. For example, aligned collagen microarchitecture is a defining feature of both fibrotic (23) and tumour ECM (24, 25), with increased alignment shown to enhance both cancer cell invasion (26) and influence fibroblast differentiation (27, 28). As such, methods to visualise and image the microarchitectural changes enable researchers to gain a better understanding of disease pathophysiology.

There are several microscopy techniques available for visualising collagen organisation both in intact tissue and cell culture. While collagen can be directly imaged without the need for extra labelling using second harmonic generation (SHG), the instrumentation required to do so (fast pulsed near-infrared illumination usually provided by a Ti:Sapphire laser) is expensive. Alternatively, for samples that are sufficiently thin to not require long wavelength imaging, collagen can be fluorescently labelled. For *in vitro* cultured samples where collagen is produced by cells, labelling is typically achieved using indirect immunofluorescence (29, 30). When collagen gels are produced synthetically, for example in studies of cultured cells interacting with ECM for investigating biological processes such as invasion of cancer cells and wound healing (31–33), the gels can be directly labelled using thiol maleimide coupling (direct conjugation of fluorophores to cysteine residues) (34) or NHS ester crosslinking (direct conjugation of fluorophores to free amine groups) (35, 36). The non-specific nature of these direct labelling methods means that they can only be performed in the absence of other proteins.

Here, we demonstrate anisotropic emission of fluorescence from labelled fibrous structures in response to excitation with polarised light. This polarised emission can bias quantitative analysis of fibre orientation of images. We have developed a method for calibrating and correcting for this bias so that orientational analysis of fluorescently labelled fibres can be performed with higher confidence.

## Materials and Methods

### Sample preparation

3D collagen matrices were created using a liquid collagen solution containing a final concentration of 2 mg/ml rat tail-derived collagen type I (Corning) with 100 μg/ml fibronectin (EMD Millipore Corp.), 2 mM HEPES (Sigma), 14.5 mM NaOH (Sigma), 0.3% NaHCO_3_ (w/v) (Sigma) and 10 μg/ml Alexa Fluor 647 NHS Ester (Invitrogen) or 10 μg/ml Alexa Fluor 555 NHS Ester (Invitrogen) made up in Opti-MEM (Gibco) supplemented with 10% FBS (v/v) (Hyclone). The liquid collagen was assembled on ice, plated in μ-Slide 4 Well chamber slides (ibidi 80426) or μ-Dish 35 mm high wall imaging dishes (Thistle Scientific 81151) and left to polymerise at 37ºC in a sterile tissue culture incubator for approximately 2 hours. Polymerised gels were kept hydrated with phosphate buffered saline (Sigma). Actin labelled with phalloidin-Alexa Fluor 488 was imaged in a pre-prepared test sample (FluoCells Prepared Slide #1, ThermoFisher, F36924). Unconjugated Alexa Fluor 647 NHS Ester dye (AF647-NHS ester) was dissolved in phosphate buffered saline to a final concentration of 0.067mg/ml.

### Polarisation-resolved TIRF microscopy

Polarisation-resolved TIRF microscopy was carried out on a customised microscope consisting of a DMi8 microscopy body and ‘SuMo’ passively stabilised stage (Leica Microsystems GmbH). Fluorescence was imaged with a 160× /1.43-NA (numerical aperture) objective (Leica Microsystems GmbH) and split into two channels by a Cairn OptoSplit system (Photometrics Dual-view/Cairn Research OptoSplit II). This was fitted with orthogonal sheet polarisers (Thorlabs) in each light path simultaneously observing parallel and perpendicular polarised fluorescence on a fast EMCCD camera (Photometrics Evolve) using a 20ms exposure (collagen and phalloidin samples). When imaging AF647-NHS ester in free solution, photobleaching was not a concern and so images were acquired as the average of 100 frames (100ms exposure). Samples were illuminated with a 638nm (Vortran) or 488nm (Dragon Laser) diode laser as appropriate at ∼ 50-100μW at the back focal plane. Laser polarisation was reduced by passing each through an optical fibre (Thorlabs). A cube polariser in the excitation path could be rotated or removed to produce vertical, horizontal, or unpolarised excitation. Each experiment was repeated on ten different fields of view (three for AF647-NHS ester in free solution) for each polarisation, producing quantitatively similar results. For each field of view the order of vertically and horizontally polarised excitation measurements was reversed to avoid any possible systemic effects of photobleaching between measurements. A ‘G factor’ image sequence (1000 frames) was recorded using ambient room light and averaged, giving the relative detection efficiencies of each channel (1.13:1). All measurements were corrected for this difference.

### SoRa microscopy

Super-resolution SoRa spinning disk microscopy images were acquired with a Nikon Eclipse Ti2 inverted microscope coupled with a Yokogawa CSU-W1 SoRa confocal scanning unit (Yokogawa Electric Company) and PVCAM Dual Prime 95B sCMOS cameras (Photometrics), utilising a CFI Plan Apochromat VC 20x objective (NA 0.75), CFI Apochromat LWD Lambda S 40XC WI (NA 1.25) and SoRa super resolution disk. Samples were excited with an excitation wavelength of 640 nm (AF647) or 561nm (AF555) from a Lighthub Ultra laser bed (Omicron-Laserage Laserprodukte GmbH) at nominal powers in the range 190-228 μW or 223-243 μW measured at the objective back focal plane, respectively. Typical camera exposure times were 60 ms. Emitted fluorescence was filtered by a quadband filter (Di01-T405/488/532/647-13×15×0.5, Semrock) and a 708/75 (AF647) or 600/52 (AF555) filter prior to detection.

### Inverted spinning disk confocal microscopy

Images were acquired using a Nikon Eclipse Ti inverted microscope with an Andor NEO SCC-01950 camera (Andor Technology) utilising a Yokogawa CSU-X1 spinning disk unit (Yokogawa Electric Company), and a Plan Apochromat VC 20x objective (NA 0.75). Samples were excited with an excitation wavelength of 638 nm from an MLC 400B laser bed (Agilent Technologies) at nominal powers in the range 414-517 μW measured at the objective focal plane. Typical camera exposure times were 100-200 ms. Emitted fluorescence was filtered using a Cy5 emission filter prior to detection.

### Polarisation measurements

Power measurements were taken using a powermeter (PM100D, Thorlabs) with a photodiode sensor (S121C, Thorlabs) attached onto a manual rotation mount along with a sheet polariser. Microscope illumination polarisation was estimated by measuring 640nm and 561nm laser power at the back focal plane with the sheet polariser rotated to different angles.

### Simulation of synthetic images

Fluorescence microscopy images of fibres were simulated in Python (Figure 5) or MATLAB (Supplementary Figure S2, Supplementary Figure S3, Supplementary Movie 1, Supplementary Movie 2). In the Python implementation, a list of 50000 fibres was created with angles *θ*_*f*_ distributed uniformly in the range [0º, 180º] and lengths randomly selected from the range [50, 250] pixels. Fibres were randomly positioned within a 5000 × 5000 pixel image and were allowed to overlap. Each fibre was assigned a base signal of 1000 counts. When polarisation effects were simulated with a polarisation angle of *θ* _*p*_, fibre signal was modified by a factor of (1 + *P* (cos^2^(*θ* _*f*_ *− θ* _*p*_) − sin^2^(*θ*_*f*_ *− θ*_*p*_))), where *P* is a factor in the range [0, 1], with higher values of *P* indicating stronger polarisation effects. The image was then blurred with a Gaussian kernel with σ = 13 pixels prior to downsampling by a factor of 5 to give an effective pixel size of 100nm. Poisson noise was then simulated, and Gaussian noise with a mean of 20 counts and standard deviation of 2 counts added. In the MATLAB implementation, the image size varied from 256 × 256 to 4096 × 512 pixels. The number of fibres was adjusted to give a consistent fibre density similar to the real data and Python implementation. The simulated point spread function was 2.7 pixels FWHM (full width at half maximum). Fibre orientations ranged from 0-180º for unaligned simulations and intrinsic alignment was simulated by restricting this range. For each simulation two images were produced: one where the fibre brightness was modified according to its orientation (polarisation operator (1 + *P* cos^2^ *θ*, where *P* is defined as above) and one where it was not, giving a ground truth for comparison. Background of 10 counts per pixels was added and Poisson noise was applied. This resulted in a typical 10:1 SNR for individual fibres. Both the degree of restricted alignment and strength of polarisation could be linearly varied across the image.

### Calculation of Fourier Ring Anisotropy spectrum

The process of calculating the emission anisotropy in Fourier space is similar to that used in Fourier Ring Correlation (FRC) (37–39). Here the steps will be laid out in a similar manner as readers familiar with FRC will find it more intuitive; those not familiar with FRC may wish to consult the suggested references. FRC estimates resolution from two independent measurements of the same structure by measuring the similarity of their Fourier transforms. For each transformed image a profile is taken around a ring, which corresponds to all coefficients of the same spatial frequency (length scale), and the linear correlation between the profile from each image is taken. Once repeated over all rings (spatial frequencies) a spectrum is produced indicating the degree of self-similarity as a function of scale. Where the correlation falls below a prescribed value (typically 1/7 for fluorescence microscopy (38)) is deemed to be the limit of resolution.

Here we wish to measure the difference between two polarisation-resolved images of the same structure rather than the size scale-dependent similarity. Specifically, we wish to know the difference in relative brightness of structural elements of different orientations. Ideally this should be done in a way that does not depend on perfect alignment of the images. With conventional fluorescence anisotropy, slight misalignment or dissimilar distortions between images would give rise to false indications of polarisation. Additionally, if we wish to measure differences as a function of angle this would mean identification and segmentation of the fore-ground structure and its orientation, which is not easy for dense and curved fibres. These issues are easily bypassed by performing the calculation in Fourier space where the dimensions of position, scale and orientation are naturally separated. A spectrum of orientation bias can then be produced in a process we name ‘Fourier Ring Anisotropy’ as follows. Conventionally in fluorescence anisotropy, where there is cylindrical symmetry around the excitation axis, the emission polarisation axes are commonly defined as being parallel and perpendicular to this axis. Here, with highly structured samples, this symmetry is not present. We therefore define emission polarisation directions in the lab frame (vertical, *V*, and horizontal, *H*). To aid the description, a graphical demonstration of the Fourier space anisotropy calculation is shown in Supplementary Movie 1.

First, the two-dimensional Fourier Transform *Y*_*pq*_ for each polarisation-resolved (horizontal and vertical) image *X*_*jk*_ is taken (example images shown in Supplementary Movie 1a, b), where the dimension of each image is *m · n*.

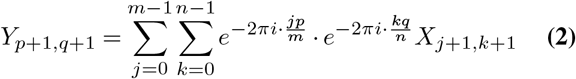

*j, k* represent image coordinates in the spatial domain, and *p, q* represent these coordinates in the frequency domain. This produces horizontal and vertical two-dimensional Fourier transforms 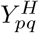 and 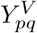 respectively. From these, a difference or ‘Fourier space anisotropy’ image *FA* is produced.

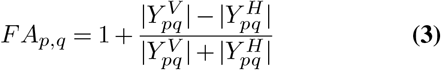

Quadrants of the *FA* image are swapped to bring the zero frequency term to the centre

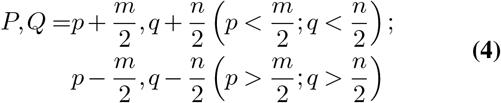

where *P, Q* and *p, q* are the new and old indices respectively (an example of a *FA* image is shown in Supplementary Movie 1c). By taking the magnitudes only of the Fourier transforms in Equation 3 all positional information of the structure, which is contained in the phase of the coefficients, is removed. The Fourier anisotropy coefficients thus only contain information on scale *r* (distance of indices from centre) and orientation *θ* (position of indices around a fixed ring of radius *r*). A coefficient *>* 1 in *FA* indicates that the structure of that scale and orientation is brighter in the vertical polarised image, whereas a value *<* 1 indicates that it is brighter in the horizontal polarised image. A set of indices corresponding to a ring *r* or (ellipse for non-rectangular images) is selected,

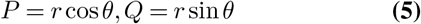

and the set of values of the selected coefficients in *F A*_*P,Q*_ drawn from it. For each specific spatial frequency *f* (corresponding to ring *r*) this sequence of values will have the form of a periodic function in *θ* with two maxima 180º apart and two minima 90º from them. These indicate the orientations with the biggest differences between the two polarisation channels. As a periodic function it can be expressed as

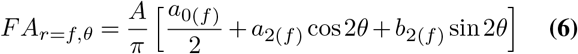

which is a standard Fourier series (Supplementary Movie 1d). The series is limited to terms of second order in *θ* because single photon selection rules limit the angular dependence of excitation probability to cos^2^ *θ*. The ‘anisotropy moments’ of this distribution can be calculated for each spatial frequency

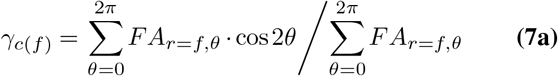

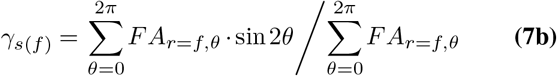

By calculating these anisotropy moments at each spatial frequency, two spectra are produced (Supplementary Movie 1e). These spectra show how polarised the fluorescence emission is as a function of scale that is independent of any intrinsic alignment of the structure as this would appear equally in both images. Additionally, the frequency-resolved output allows the differentiation of signals with different degrees of polarisation, such as the background (which should display low polarisation from out-of-focus signal at low frequencies) and noise (unpolarised, at high frequencies).

### Fourier ring depolarisation (FouRD)

The process for deriving a spectrum and performing a correction is very similar to that for obtaining the Fourier anisotropy spectrum from a pair of polarisation-resolved images (as described in “Calculation of Fourier Ring Anisotropy spectrum”), but only a single image is used where polarisation is not resolved. In this case, the absolute of the Fourier transform of this image, *Y* ^*u*^, contains orientation information on both the polarisation bias and the intrinsic fibre alignment. Accompanying the following description, a graphical representation of the process is shown in Supplementary Movie 2. To use just a single image, Equation 3 above is replaced by:

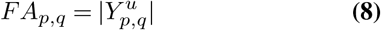

The cosine and sine moments *γ*_*c*_, *γ*_*s*_ can then be calculated as before for this image. Even without polarisation-resolved detection, the polarisation bias of the system can be determined using this method. This is because in Fourier space the observed image *Y* ^*ob*^ can be expressed as the true image *Y* ^*T*^ modified by an excitation operator *Y* ^*ex*^.

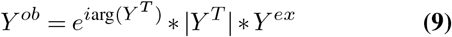

*Y* ^*ex*^ describes the angular variation in excitation probability at each spatial frequency. This operator, being real, only modifies the amplitudes of the Fourier-transformed image (brightness in real space), leaving the phase (first term in Equation 9 describing position/structure in real space) unchanged.

If a calibration sample is imaged, which is similar to the one under examination (fluorophore, labelling method, structure type being labelled, and imaging conditions) except for having no intrinsic structural alignment, the resulting spectra will be affected only by polarisation. So long as the image contains a large number of randomly orientated fibres, | *Y* ^*T*^| will have negligible second order angular dependence in *θ* (*γ*_*c*_, *γ*_*s*_ = 0). Measuring the moments of |*Y* ^*ob*^| will thus only be affected by *Y* ^*ex*^. This fully describes *Y* ^*ex*^ as it is itself limited to terms of second order in *θ* (see above). The resulting spectra of the calibration sample describe how each orientation at each spatial frequency is over-or under-represented in the image as a result of the laser illumination polarisation (Supplementary Movie 2 a-e).

To remove polarisation bias from an image, an inverse of the above-described excitation operator can be constructed. This is possible because, for single-photon excitation, most fluorophores only have a very small angle between their absorption and emission dipole moments (2). Multiplying (direct product) the Fourier transform of the observed image by this operator *Y* ^*inv*^ removes the effect of the polarised excitation, producing a corrected image *Y* ^*c*^

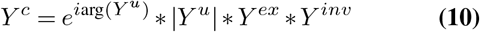

The last two terms cancel each other, leaving *Y* ^*c*^ as a true representation of *Y* ^*T*^. The inverse operator can be produced from the spectra derived from the calibration sample. For each spatial frequency the Fourier ring profile is calculated from the respective moments

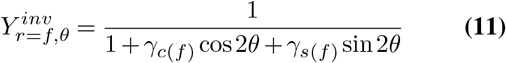

The inverse operator is constructed by combining all of these ring profiles to form a Fourier space image. The evolution of this inverse operator for each radius is shown in Supplementary Movie 2e, with the corresponding Fourier image building up in Supplementary Movie 2f. Combining all the ‘rings’ for each frequency *r* = *f* and then swapping quadrants as before (reverse of Equation 4) produces the Fourier space inverse operator image.

The excitation operator *Y* ^*ex*^ is a joint property of the microscope, the fluorescent label and its environment. However, it is independent of the structure, and thus should be common to all measurements performed on the same system under similar conditions. Therefore it should be possible to correct all such measurements using the same inverse operator. As *Y* ^*inv*^ only cancels the effects of *Y* ^*ex*^, any intrinsic orientation dependence of *Y* ^*u*^ (fibre alignment) is left unmodified. To correct an image *X*_*u*_, its Fourier transform is multiplied (direct product) by this operator and the inverse transform taken

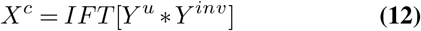

where *X*^*c*^ denotes the corrected image in real space, and *IFT* represents the inverse Fourier transform. Supplementary Movie 2g shows an example of *Y* ^*u*^ *\*Y* ^*inv*^ prior to swapping the quadrants, and Supplementary Movie 2h shows the resulting *X*^*c*^ image produced from using this. As described above, the inverse operator, being real, only modifies the absolute values of the transformed image; the structure (phase in Fourier space) of the original real space image *X*^*u*^ is un-modified, but the relative brightness of features at different orientations is changed. The advantage of the inverse operator being a function of frequency is that larger and smaller scale elements of the image (background and noise respectively) that have no initial bias do not have bias artificially introduced. As an optional step, very noisy spectra can be smoothed by a method such as n-point moving average. So long as the smoothing is insufficient to change the overall shape of the spectrum, it will have minimal effect on the output image. This is demonstrated in Figure 6g). No smoothing was used for any other corrections.

The above process assumes that the excitation polarisation is a constant across the field of view. Where this is not the case, the calibration image and the image to be corrected can be divided into smaller image patches and the calibration/correction procedure applied individually to each patch and then recombined. This was the case for all images taken on the SoRa microscope. The patches must be small enough that the polarisation is near constant across the patch, but still large enough to contain a representative distribution of fibre orientations. When recombining the full image, slight low frequency artefacts can be notices at the patch boundaries. These are due to discontinuities in frequency space, particularly in the lower frequency components, arising from different corrections being applied to different areas of the image. Such discontinuities can be easily avoided by starting the correction at a slightly higher spatial frequency and leaving the lowest frequencies unmodified. By default the starting frequency is *f* =2 when doing the patch-wise correction, which is normally adequate. As demonstrated in the Results section the choice of start frequency has negligible effect on the AFT analysis of fibre alignment.

### Image acquisition, processing and analysis

Images on the SoRA system were acquired using NIS-Elements AR Version 5.42.02, whilst images on the inverted spinning disk were acquired using NIS-Elements AR Version 5.21.03. Polarisation-resolved images were acquired using Micro-Manager Version 1.4. Postprocessing was done in Fiji (40). *AFT analysis:* Alignment of fibres in real and simulated data were quantified in the MATLAB implementation of Alignment by Fourier Transform (AFT). 250 pixel windows with a 50% overlap were used throughout; corresponding calibrated window sizes are stated in figure captions. Neighbourhood radius was set to 2 by default, although the calculated order parameter was not used for analysis in this work. No masking or thresholding was applied prior to performing analysis. AFT output was modified so that in addition to outputting angle maps, eccentricity maps were also generated. These have values from 0-1 where higher eccentricity indicates a higher degree of anisotropy in the analysed patch. *OrientationJ analysis:* OrientationJ version 2.0.4 was used for analysis, and outputs displayed here are the ‘orientation’ output and histograms of this image. For simulated data (Figure 5a, b and Supplementary Figure S3c), local window *σ* =2 was used. For 40 × data (Figure 5c) *σ* = 4.4, and for 20 × data (Figure 5d) *σ* =3 was used. *CytoSpectre analysis:* CytoSpectre version 1.2 was used for analysis, with all parameters set to default values except for magnification and pixel size, which were set according to the image being analysed. Displayed outputs are the ‘mixed component’ spectrum.

## Results

### Fluorescently-labelled collagen fibres emit polarised fluorescence

Collagen fibres labelled with Alexa Fluor 647 NHS ester (henceforth collagen-AF647; dye and linker structures shown in Figure 1c and Figure 1d) were imaged in a TIRF microscope with polarisation-resolved detection. Fluorescence is split with a 50:50 beam splitter, sent through orthogonal polarisers and imaged ‘side by side’ on the camera chip. Where a horizontal polarisation filter is situated in the detection pathway, fibres orientated horizontally in the image plane appear bright relative to those that are vertically orientated (Figure 2a, green arrows). Similarly, in the channel with a vertical polarisation filter, vertically orientated fibres appear bright relative to the horizontally orientated fibres (Figure 2b, magenta arrow). These differences can be seen clearly when the two images are overlaid in contrasting colour channels (Figure 2c).

**Fig. 2.**
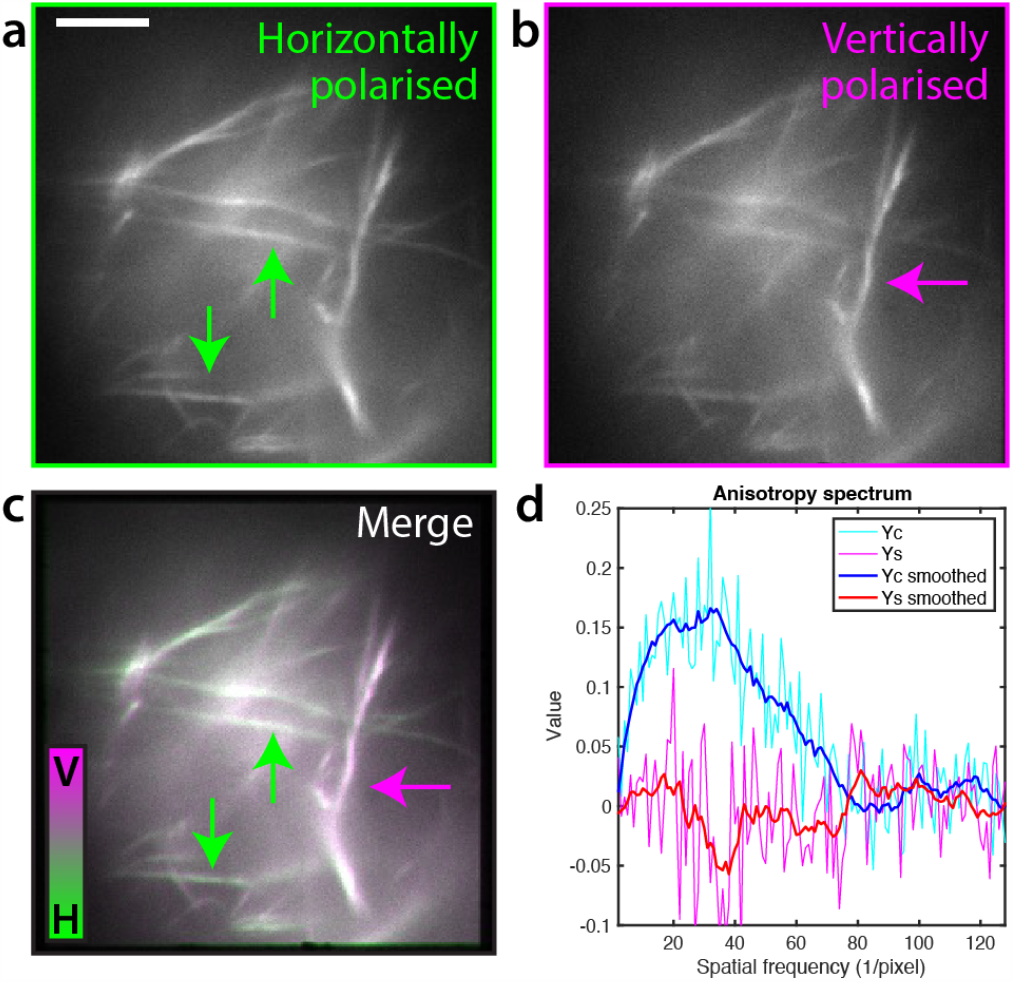
Polarisation-resolved detection reveals polarised emission correlated with fibre orientation in collagen-AF647. **a** Horizontally and **b** vertically polarised fluorescence images of collagen gels comprising collagen monomers labelled with Alexa Fluor 647 via NHS ester linkage. Arrows indicate examples of bright fibres parallel to the polarisation direction. **c** Overlay of the horizontally polarised image from **a** (green) and vertically polarised image from **b** (magenta). The colour ratio clearly varies with fibre angle. **d** Quantification of the polarisation bias can be plotted in the Fourier anisotropy spectrum. Light blue and pink lines show the raw *γ* _*c*_ and *γ* _*s*_ components respectively; dark blue and red lines are smoothed with a 3-point moving average. Scale bar = 5 μm.

Quantification of image anisotropy is not necessarily straight-forward; for existing methods controls must be done to account for differences in detection efficiency in the two polarisation channels (G factor calculation) (2). If the fluorescent signal is split according to polarisation and imaged simultaneously as two separate channels, the images must be perfectly aligned and any differences in magnification and distortion accounted for and corrected. If instead separate polarisation-resolved images are acquired sequentially, there may be selective photobleaching of fluorophores (orientational hole burning) between measurements. Either of these could give rise to false indications of polarised emission. Additionally, there may also be differences in polarisation between fluorophores immobilised on structures (here, collagen fibres), out-of-focus fluorescence, and free fluorophore diffusing in the background, which would require segmentation of the structure for analysis. Therefore, we developed a method for analysing polarisation effects in Fourier space, which allows for inspection of polarisation at different spatial frequencies (length scales) within the image that is robust to spatial misalignment/distortion between channels (Fourier Ring Anisotropy, Methods). Figure 2d shows how the strengths of cos 2 *θ* (*γ* _*c*_, blue) and sin 2*θ* (*γ* _*s*_, red) polarisation components vary with spatial frequency for the field of view imaged with polarisation-resolved microscopy. The cos 2*θ* component is substantially positive at intermediate spatial frequencies (∼5 - 55 units (1/pixel), corresponding to length scales of 5 - 0.5 μm), whereas it is close to 0 for very high frequencies (noise) and very low frequencies (background fluorescence). This quantitatively confirms the qualitative observation from the polarisation-resolved images that emitted fluorescence is polarised parallel to the collagen fibres. Meanwhile, the lack of any significant sin 2*θ* component across all spatial frequencies is the expected result of polarised detection with polarisers crossed at 90º and this choice of axis. It does however indicate the scale of noise effects in the calculation and therefore the significance of the cos 2θ result. Moreover, the measured polarisation components are similar regardless of the excitation polarisation (no photoselection), suggesting highly orientationally constrained fluorophores. Therefore, these results indicate that collagen-AF647 exhibits polarised fluorescence parallel to the orientation of the fibres.

We next investigated whether this phenomenon was a specific property of collagen-AF647, or whether this effect was a combination of biological structure, linker molecule and fluorophore identity. Collagen gels labelled with Alexa Fluor 555 NHS Ester (collagen-AF555; dye structure shown in Figure 1c) also showed polarised emission, albeit to a lesser extent than collagen-AF647 (Figure 3a, 3b), indicating that the structure of the fluorophore is a likely contributor to the underlying mechanism of this orientational restriction. Polarisation-resolved imaging and Fourier domain analysis of fixed cells with the filament-forming cytoskeletal component F-actin labelled with Alexa Fluor 488 phalloidin (actin-AF488; dye and linker structures shown in Figure 1c and Figure 1d) displayed even stronger polarisation effects than collagen-AF647 (Figure 3c, 3d). Again, the direction of the polarised emission is independent of the excitation polarisation direction, indicating fluorophore are constrained to a narrow range of angles relative to the fibre they are attached to. Meanwhile, a control image taken of AF647-NHS ester in aqueous solution, where fluorophores are unconstrained and undergo free rotational diffusion on a timescale shorter than their fluorescence lifetime (*τ*_rot_ ≪ *τ*_*f*_, Equation 1), displays no visible qualitative difference between the two polarisation channels, and both its *γ*_*c*_ and *γ*_*s*_ components are constant about zero across all frequencies (Figure 3e, 3f). Taken together, these results indicate that the polarisation of emitted fluorescence from labelled fibrillar biological structures is dependent on a wide range of factors.

**Fig. 3.**
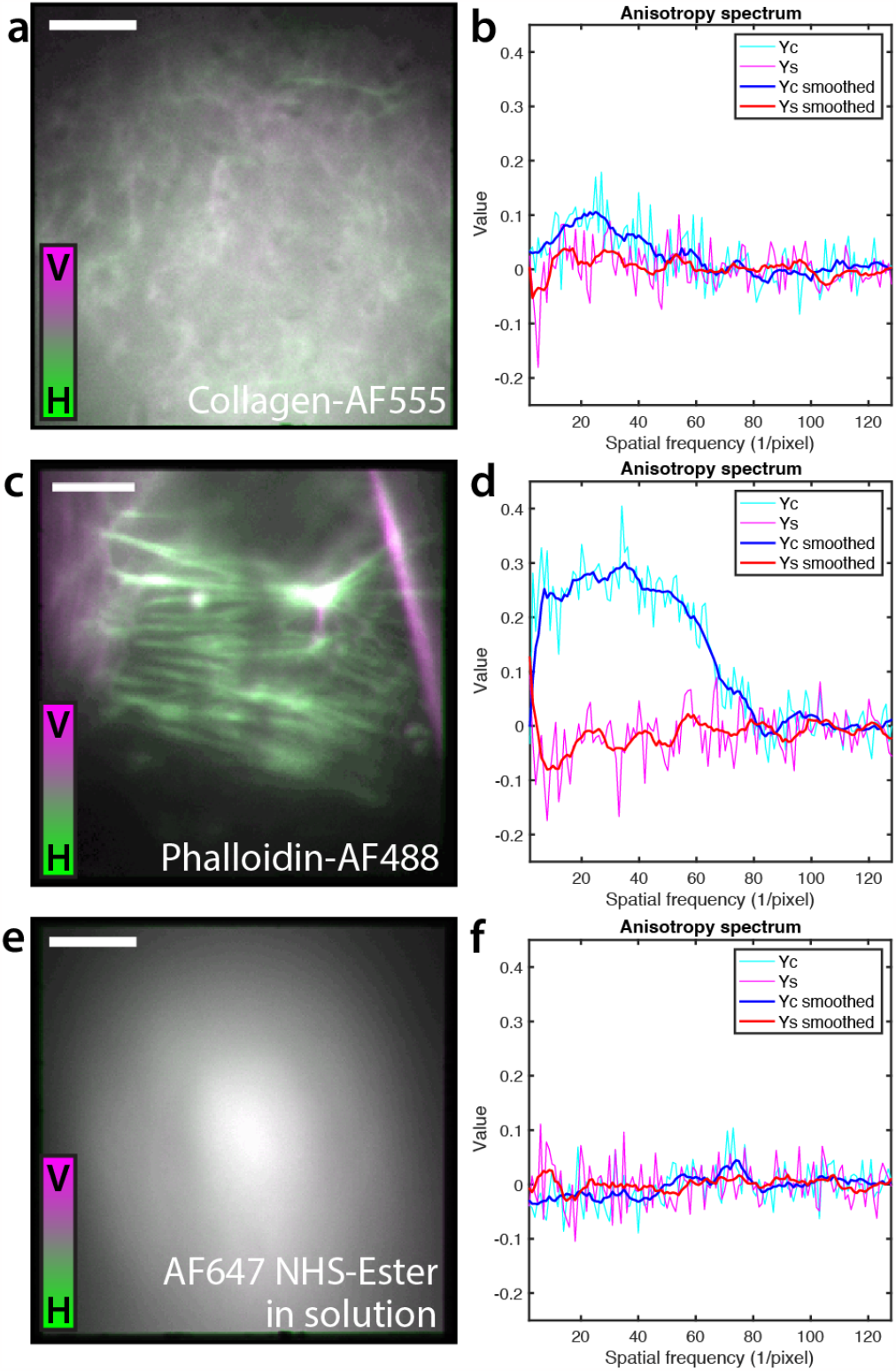
Different combinations of structure and labelling show different degrees of polarised emission. **a** Overlaid horizontally (magenta) and vertically (green) polarised fluorescence images of collagen-AF555. **b** Fourier anisotropy spectrum of collagen-AF555 with *γ*_*c*_ and *γ*_*s*_ components in pink/red and light blue/dark blue respectively. **c** Overlaid horizontally (magenta) and vertically (green) polarised fluorescence images of actin labelled with Alexa Fluor 488 phalloidin (actin-AF488). **d** Fourier anisotropy spectrum of actin-AF488. **e** Overlaid horizontally (magenta) and vertically (green) polarised fluorescence images of Alexa Fluor 647-NHS ester in aqueous solution. **f** Fourier anisotropy spectrum of aqueous Alexa Fluor 647-NHS ester. The measured anisotropy of the bound systems vary greatly, whereas as expected the dye in free aqueous solution shows no fluorescence anisotropy. Scale bar = 5 μm.

### Polarisation effects can mask alignment of collagen fibres

While for some applications it is useful to study small fields of view of fibres at high resolution as in Figures 2 and 3, many biological studies are more concerned with alignment and characteristics of fibre matrices on a larger scale (41, 42). We therefore moved to potential polarisation effects on larger fields of view of collagen-AF647 with spinning disk confocal microscopes.

In the absence of any biological stimuli, for example cells seeded within the the matrix, or physical biases such as tilting the gel during polymerisation, fluorescently-labelled collagen is expected to polymerise randomly with all fibre orientations represented equally. However, upon analysing spinning disk confocal images of large field of view images with AFT, strong quantitative angle biases were observed that did not qualitatively agree with local fibre organisation (Figure 4). Indeed, as shown in Figure 4a, the vectors indicating AFT-calculated local directionality of the matrix showed a large-scale elliptical pattern across the whole field of view. When local patches of this image were inspected, it appeared that fibres which were subjectively assessed to display strong local alignment were not being represented in the objectively calculated angles (Figure 4b). Similar biases were picked up when the data was analysed using OrientationJ and CytoSpectre (Figure 5). For simulated randomly oriented fibres with no polarisation bias (Figure 5a), the AFT and OrientationJ angles maps showed a random distribution of angles, and the CytoSpectre orientation distribution plot appeared uniform with a high circular variance (0.99) indicating near-perfect isotropy. Adding a polarisation operator (20% contribution of excitation aligned to 80º) to this simulated distribution resulted in an enrichment of angles close to 80º in the AFT and OrientationJ angle maps, and a decrease in the circular variance of the CytoSpectre plot to 0.93. The CytoSpectre orientation plot also became biased towards a mean orientation of 169º (the direction of 0º was 90º offset from AFT and OrientationJ) (Figure 5b). Real collagen-AF647 gels imaged at 40× magnification (Figure 5c) and 20× magnification (Figure 5d) showed strong directional biases in the AFT and OrientationJ angle maps. Similarly, the CytoSpectre plots displayed low circular variance (0.893 at 40× and 0.894 at 20×) indicating anisotropy within the image, with mean orientations of 0.1º and 4.4º (corresponding approximately to the mean angle measured in the AFT and OrientationJ analyses when reported in the 90º-shifted frame of reference). These results suggest that the apparent strong alignment reported by AFT in the samples which should be randomly arranged is not a bias in analysis, as the same alignment also manifests when the analysis is performed on a more local scale using OrientationJ and globally across the image using CytoSpectre.

**Fig. 4.**
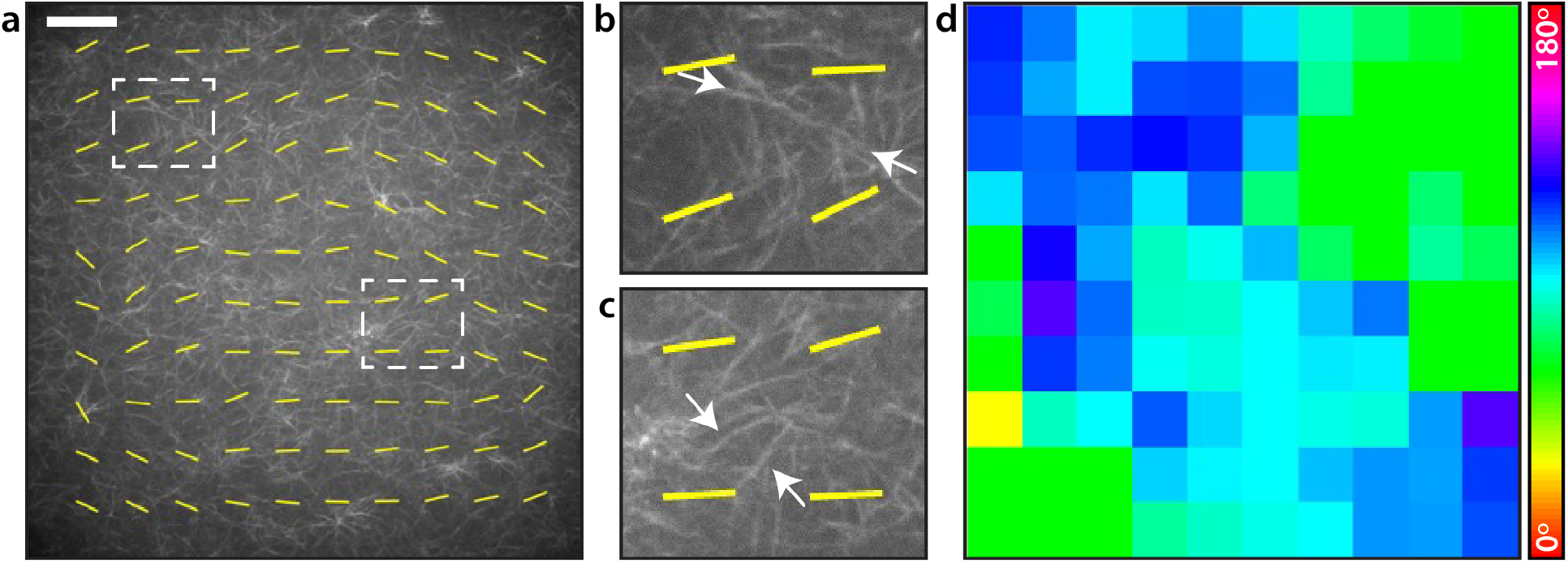
Polarisation effects dominate orientational analysis of randomly polymerised collagen AF-647 gels. **a** Image of a single optical slice of unaligned collagen-AF647 matrix imaged at 40 × magnification on the SoRa microscope. Yellow overlay indicates alignment vectors generated by AFT analysis. White dotted boxes indicate regions magnified in panels **b** and **c. b, c** Magnified insets showing local structure of the collagen matrix and calculated AFT alignment vectors. White arrows indicate bright fibres which would be expected to influence local alignment calculations, but which do not agree with generated vectors. **d** Angle heatmap generated by AFT analysis with the local angle colour coded according to the colour bar. The predominant angle in the image was 89 ± 20º (mean ± standard deviation). AFT analysis was performed with 250 pixel window size, corresponding to 71.5 μm. Scale bar = 50 μm.

**Fig. 5.**
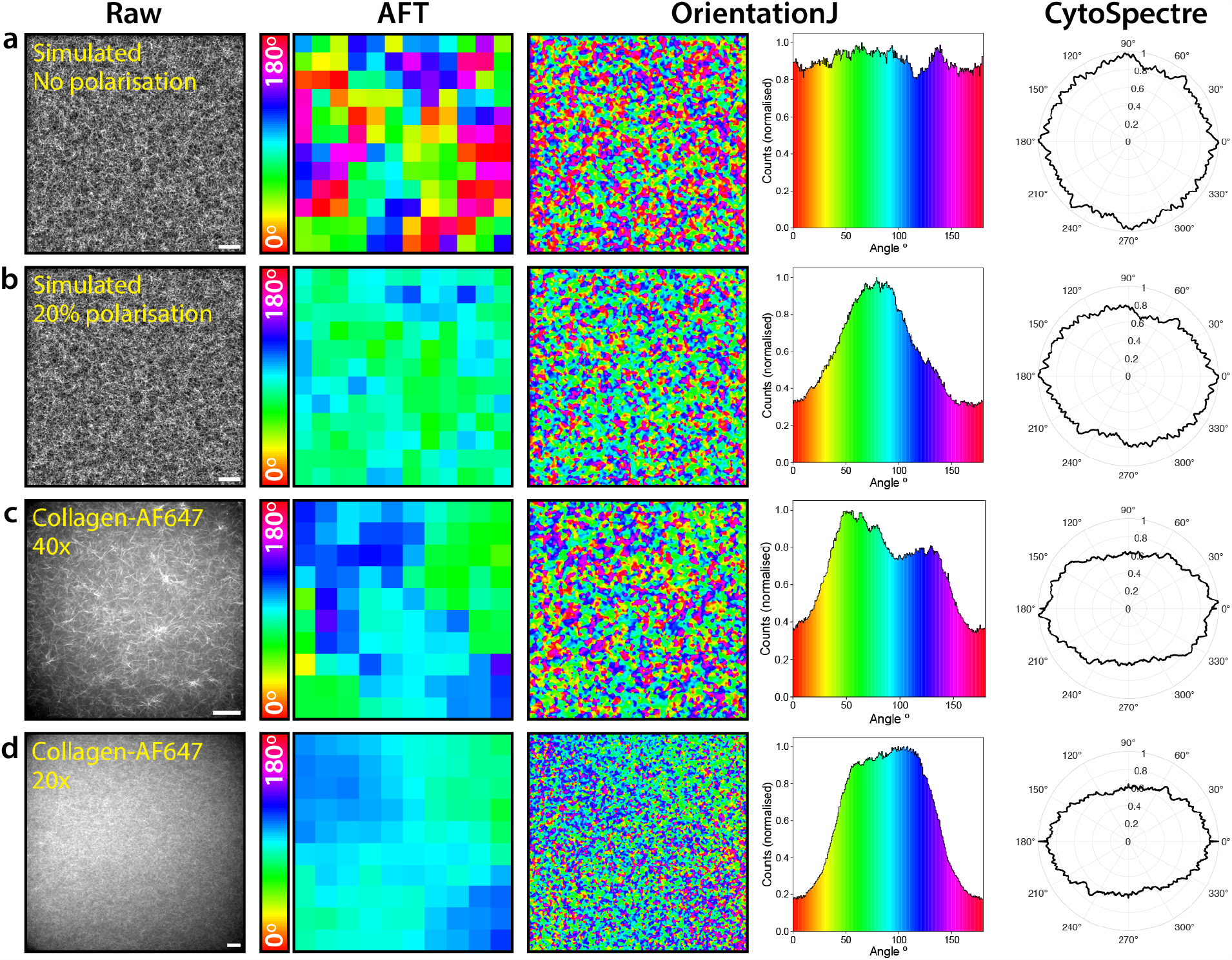
Polarisation bias affects different orientational analysis algorithms for both simulated and real data. Simulated and real data were analysed with three different open source fibre alignment algorithms: AFT, OrientationJ, and CytoSpectre (analysis parameters listed in Methods). The ‘Orientation’ colourmap output of OrientationJ is displayed alongside a histogram of this colourmap. Colour bar for AFT and OrientationJ angle maps is the same. CytoSpectre output represents the ‘mixed’ component. Images and analysis are shown for **a** simulated randomly oriented fibres with no polarisation effects, **b** the same simulated fibres as in **a** but with 20% polarisation at an angle of 80 °, **c** collagen-AF647 gel imaged on SoRa spinning disk confocal microscope at 40× magnification, **d** collagen-AF647 imaged on SoRa spinning disk confocal microscope at 20× magnification (note: different sample and day to image acquired in **c**). Scale bars: **a, b** = 10 μm, **c, d** = 50 μm. AFT block size was 250 pixels for all analysis, corresponding to 25 μm in **a** and **b**, 71.5 μm in **c**, and 143 μm in **d**

**Fig. 6.**
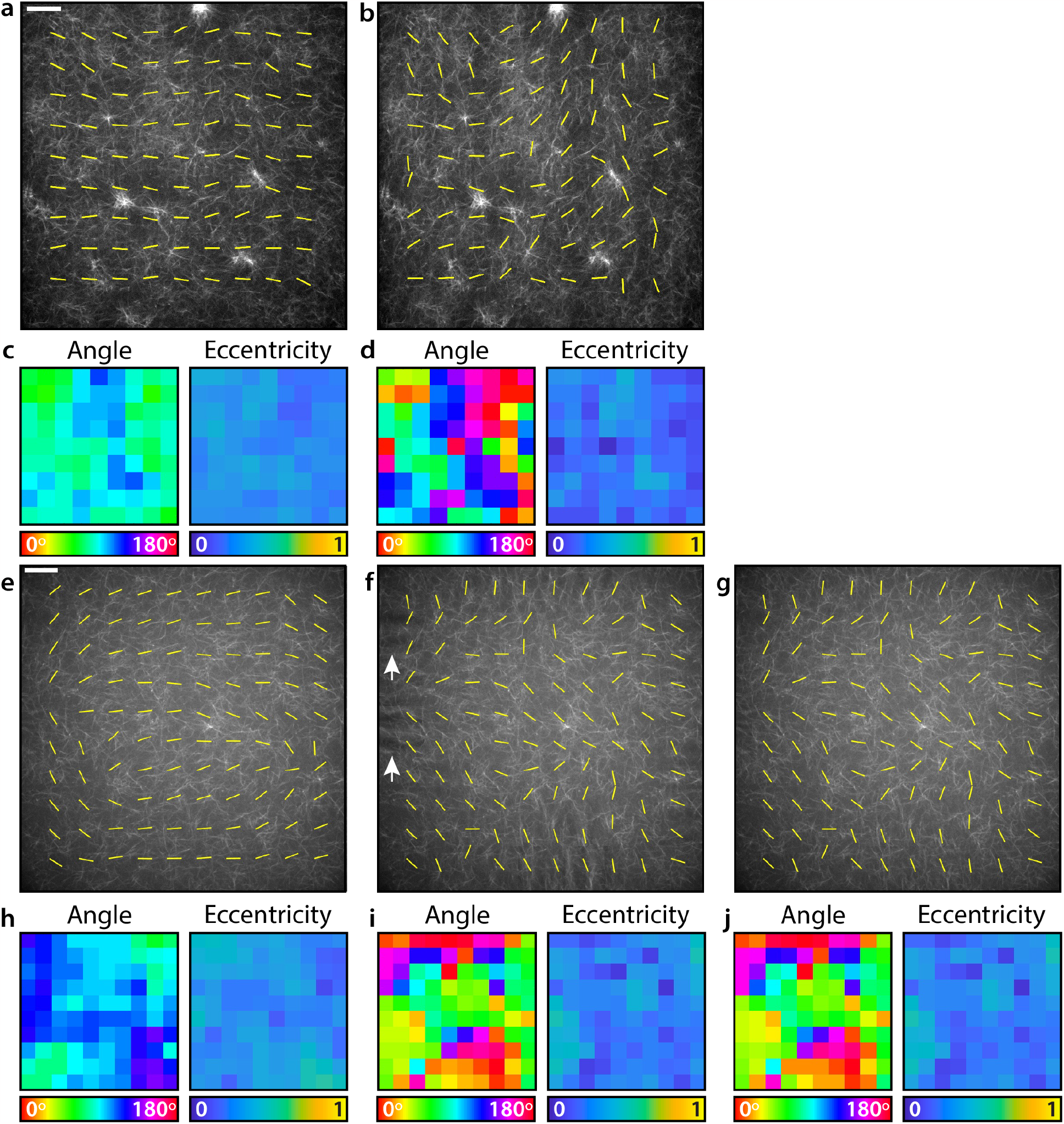
FouRD correction of collagen-AF647 gels acquired using spinning disk confocal microscopes. **a** Image of a collagen-AF647 gel acquired on an inverted spinning disk confocal microscope. Yellow lines indicate the vector map generated by AFT. Scale bar = 40 μm. **b** The result of applying FouRD correction to the image in **a. c** The angle map (which shows the same information as the yellow vector lines, but as a colour plot) showing global orientational bias, and eccentricity values from analysing the image in **a** with AFT. **d** The angle map and eccentricity values from analysing the FouRD-corrected image in **b** with AFT. **e** Image of a collagen-AF647 gel taken on a SoRa spinning disk confocal microscope. Scale bar = 40 μm. **f** The result of applying FouRD correction to the image in **e**, using rings starting at radius = 2 and without smoothing the spectrum. White arrows indicate artefacts introduced at image edges. **g** Applying FouRD to the image in **e**, but starting from radius = 5 and smoothing the spectrum prior to inversion and application, does not alter the fine structure in the image but removes the edge artefacts. **h** Angle and eccentricity maps from AFT analysis of the image in **e. i** Angle and eccentricity maps from AFT analysis of the FouRD-corrected image in **f. j** Angle and eccentricity maps from AFT analysis following modified FouRD correction to prevent image artefacts show no appreciable difference from the maps in **i**. On both microscopes, the strong orientational bias indicated by AFT is completely removed by FouRD leaving the expected random distribution of an unaligned sample. AFT block size for all analysis was 250 pixels, corresponding to 72 μm in **a**-**d** and 71.5 μm in **e**-**j**

To confirm whether this quantitative orientational bias was due to polarisation effects in collagen-AF647, we rotated the chamber containing the collagen matrix by ∼ 90º and performed AFT analysis on the images acquired before (Supplementary Figure S1a) and after this rotation (Supplementary Figure S1b). If the alignment quantified by AFT analysis were due to real directional organisation of collagen fibres within the matrix that we were unable to perceive subjectively, then the expectation is that physical rotation of the sample would result in a corresponding shift in the measured angle distribution. However, the same angle was dominant in the angle analysis before and after sample rotation. The same rotational invariance was reported when repeating the experiment on collagen-AF555 (Supplementary Figure S1c, Supplementary Figure S1d). For the polarisation-resolved experiments in Figure 2 and Figure 3, we had experimental control over the excitation polarisation; however, on the commercial spinning disk confocal system, we did not have such control over the input polarisation, which was provided via a single mode polarisation maintaining optical fibre from the laser bed. As a result of our findings in the polarisation-resolved system and the sample rotation measurement, we sought to assess polarisation in the spinning disk system by measuring the laser power at the back focal plane of the objective after a sheet polariser which was incrementally rotated. The sinusoidal variation in intensity in response to polariser angle confirmed a substantial degree of polarisation in the excitation lasers (Supplementary Figure S1e, S1f).

A key question is: to what extent can orientation arising from polarisation effects mask the true alignment of fibrillar structures in a sample? To investigate this, we simulated populations of fibres with a range of intrinsic alignments and subject to a range of polarisation strengths (Supplementary Figure S2). We first simulated a population of fibres subject to 50% vertically polarised excitation light (with the remaining 50% unpolarised) and increased the net alignment of the fibres from uniformly random over 180º to horizontally constrained over a range of 150º (Supplementary Figure S2a). For randomly oriented fibres subject to 50% vertically polarised excitation, AFT analysis outputted vertically aligned vectors. As the fibre population became more intrinsically aligned, a midpoint was reached where the competing effects of polarisation and intrinsic alignment cancelled each other out to give a seemingly random orientation in the AFT output. Eventually a degree of fibre alignment was reached which out-competed the effect of the polarisation, and the AFT angle distribution matched the input simulated fibre alignment. Similarly, a net horizontally aligned fibre population was simulated, but this time subject to excitation ranging from un-polarised to 50% vertically polarised (Supplementary Figure S2b). AFT analysis of this data showed that at low excitation polarisations the calculated vectors accurately reported the known fibre alignment. However, again, as polarisation effects became stronger these cancelled out and eventually dominated the structural alignment. As a result, the effective angles reported by AFT were representative of the polarisation effects in the system rather than the true alignment of the fibres. A notable observation from these simulations is that it is very difficult to visually assess the difference between ‘alignment’ introduced by polarisation (i.e. where fibres parallel to the polarisation direction appear more intense than those perpendicular to the polarisation) and true fibre alignment.

### Polarisation effects can be corrected on the image level using Fourier Ring Depolarisation

The simulation results in Supplementary Figure S2 demonstrate that if structure is already substantially aligned in the sample and the polarisation is weak, then the effects of polarisation do not impact automated quantification. However, if a study requires the quantification of subtle alignment effects, for example in a scenario where the researcher wishes to establish an experimental timepoint at which matrix organisation transforms from random to orientated, polarisation effects can significantly disrupt quantification. To mitigate for this, we developed a method to correct for polarisation effects called Fourier Ring Depolarisation (FouRD).

FouRD requires the acquisition of a calibration image of the same labelled structure on the same microscope as the data to be corrected. The calibration sample should contain the structure of interest labelled identically to that in the sample to be corrected, in the same type of imaging chamber, and should be assumed to have a random orientational distribution. This image, having no intrinsic alignment, will only contain information on the polarisation effects of the system. The calibration image is first analysed using a very similar Fourier analysis as for Figure 2d, but with only a single non polarisation-resolved image required (Methods; Equations 8 and 9) to calculate its anisotropy spectrum. This is demonstrated for simulated data of randomly distributed fibres with an excitation operator of 1 +cos^2^(θ + 45 °) in Supplementary Figure S3a, S3e. For demonstration, a test image was simulated which had the same polarisation operator as the calibration image (i.e the same ‘microscope’) but had intrinsically horizontally aligned fibres within a ±75º distribution (Supplementary Figure S3b, S3f). Using the calibration polarisation spectrum in Supplementary Figure S3e, an inverse polarisation operator was calculated and applied to the test image. The inverse operator contains the opposing orientational bias to the calibration image. Applying this operator to the image (multiplication in Fourier space) effectively cancels the effects of polarisation. The results of applying this correction to the image were assessed via AFT analysis (Supplementary Figure S3c) and examination of the Fourier anisotropy spectrum (Supplementary Figure S3g). Both methods showed excellent agreement with simulated ground truth data which had an identical fibre distribution to the test image but without the polarisation effects (Supplementary Figure S3d, S3h). When the corrected and ground truth data were analysed using OrientationJ (S3i) and CytoSpectre (S3j), both these methods also reported excellent agreement between the two images and a substantial different from the uncorrected image.

The efficacy of FouRD in correcting the effects of polarisation bias in fibre alignment quantification for real collagen-AF647 matrices acquired on two different spinning disk confocal microscopes is demonstrated in Figure 6. In the first spinning disk system, FouRD correction removes a dominant angle bias from randomly oriented fibres (Figure 6a-d), changing the distribution of angles measured using AFT from 84 ± 16º to 97 ± 48º (mean ± standard deviation), a distribution where all angles are essentially equally likely. While FouRD changed the locally calculated direction vectors, it does not reduce the eccentricity associated with each of these vectors to zero (0.32 ± 0.05 before correction compared to 0.24 ± 0.08 after correction). Eccentricity is an indicator for the ‘strength’ of the orientational effect; values close to 1 indicate a higher degree of anisotropy at the reported angle, whereas values closer to 0 indicate that the patch is more isotropic. The low values reported after correction probably reflect the local degree of alignment within the area and the contribution of insufficient number of fibres and noise. As the eccentricities can only be positive, a small positive mean would be expected for an unaligned sample. In the SoRa spinning disk system, the AFT results on the uncorrected image (Figure 6e) indicate the laser polarisation is not uniform but has significant spatial variation. In this case the calibration and test images were broken into 7 × 7 tiled images, and the correction applied individually to each tile (see Methods). One of the benefits of the FouRD method is that the frequencies that is applied over can be limited. With a tiled correction, if all frequencies are used for correction, then the edges and/or tile boundaries of the corrected image can display some visible artefacts (Figure 6f, white arrows). These arise from discontinuities at the lowest spatial frequencies. Such artefacts can be avoided by instead starting the FouRD correction from a slightly higher spatial frequency; Figure 6g shows the same image corrected using the same anisotropy spectrum but smoothed with a 3 point moving average and staring from a ring of radius 5 (as opposed to a radius of 2 for Figure 6f). Importantly, however, changing the starting point of the correction on this frequency scale has no noticeable impact on the resulting AFT analysis (Figure 6i, 6j), indicating the robustness of the FouRD technique.

Furthermore, Supplementary Figure S4 demonstrates the performance of FouRD correction when using the same calibration image to correct six different images of randomly oriented collagen-AF647 acquired on the SoRa system. In all cases, FouRD correction greatly reduces the prominence of the dominant angle retrieved by AFT analysis, and shifts the location of this peak to a random location indicating both that no polarisation effects remain in the image and that no new angular bias is written in to the data as a result of FouRD correction.

## Discussion

Although polarisation effects are well-known by researchers within the fields of optics, spectroscopy and specialised fluorescence applications such as FRET or FLIM, researchers using commercial or facility fluorescence microscopy systems may be less aware of them. Indeed, many routinely used confocal microscopes are likely to have polarised illumination due to the use of lasers (which often emit linearly polarised light) and delivery to microscope bodies via polarisation maintaining optical fibres. Angled optical components in the beam path (such as the major dichroic) can even induce small polarisations to an unpolarised source. To complicate matters further, on-sample illumination polarisation will also vary according to the laser line and the objective used for imaging; for example, objectives with a lower numerical aperture will typically preserve polarisation better than ones with higher numerical apertures (43).

For many researchers this may be of little consequence or interest. For example, for biological researchers wishing to make subjective observations of quantitative measurements unrelated to directionality, polarisation effects can likely be safely ignored. However, as shown here, polarisation effects can impact quantification of narrow fibrillar structures such as collagen fibres. In recent years there has been an acceptance that cultured cells in 2D monolayers likely behave differently to cells within 3D environments (44). As a result, an increasing number of *in vitro* studies are moving towards spheroid and organoid assays within 3D matrices that better approximate native extracellular matrix as a compromise between using cell culture models and animal models (45, 46). In addition to enabling study of cell-cell interactions in 3D, moving away from 2D monolayers also provides more opportunity to examine cell-matrix interactions. Manual analysis of local organisation of fibrillar matrix proteins such as collagen is impractical and unreliable, and so studying aspects of cell-matrix interactions relies upon automated computational methods such as AFT, OrientationJ and similar. Therefore, it is critical to report and understand any imaging properties that may bias measurements from these methods.

Here, we observed that collagen-AF647 exhibits polarised emission in three different microscope systems. Polarisation-resolved detection on the TIRF system indicated polarised emission parallel to fibre axes, while detection without polarisation separation on two commercial spinning disk confocal microscopes with polarised excitation generated images where polarised emission dominated quantitative orientation analysis. The common assumption in fluorescence microscopy is that when a structure is labelled with fluorophores, these are either static in random orientations or can rotate on a timescale that is fast relative to their fluorescence lifetimes (Figure 2a, Equation 1). However, our observations here suggest that a substantial proportion of the fluorophore population can be restricted to a narrow distribution of orientations (Figure 1b). The mechanism underlying this behaviour is multifactorial. Firstly, it appears that for collagen the identity of the fluorophore is one determinant of orientational restriction (and hence polarisation). This is somewhat surprising, as Alexa Fluor 647 and Alexa Fluor 555 have very similar structures, differing only by one methylene group within the cyanine core (Figure 1c). However, previous studies have shown that Alexa Fluor 555 has a faster rotational lifetime than Alexa Fluor 647, which may contribute here to the lower anisotropy observed between collagen-AF647 and collagen-AF555 (5). As Alexa Fluor 555 and Alexa Fluor 647 are both cyanine dyes, we attempted to make a collagen gel labelled with Alexa Fluor 488, which is from the structurally distinct xanthene dye family (Figure 1c). However, collagen-AF488 failed to correctly polymerise and instead formed a gel with diffuse fluorescence and only a small number of well-defined fibres. This suggests that the identity of the fluorophore is an important consideration when labelling collagen gels for polymerisation. Our observation that actin-AF488 also displays polarised emission suggests that this phenomenon is not specific to cyanine dyes and collagen. This is in agreement with previous work where actin filaments were labelled with a range of different fluorophores with the linker placing the fluorophore at various residues and binding sites, and fluorescence polarisation measured using a microspectrophotometer (rather than imaged) (47). In this study, FITC-phalloidin was shown to label actin with its dipole aligned parallel to the fibre axis (FITC is also a xanthene derived dye, like Alexa Fluor 488). However, some labelling methods, for example the dye dansyl ethylenediamine enzymatically bound to a specific residue, resulted in dipole orientation perpendicular to the actin fibre axis. While fluorophore alignment with some biological fibres has been described before, this has not been in the context of imaging and quantifying fibre alignment. Here we have not investigated the role of the linker molecule connecting the target protein and the fluorophore. One would expect that a larger linker molecule would have different rotational properties to a smaller linker molecule (e.g. phalloidin compared to NHS ester, Figure 1d). One avenue for future research is to investigate the effect of different sized linkers, such as NHS esters, primary immunofluorescence and secondary immunofluorescence, on fluorescence polarisation from the same fibrillar protein.

If polarisation effects are expected within a sample, the only definitive method for confirming this is using a polarisation-resolved system as for the data in Figure 2 and Supplementary Figure 3. However, if this is not possible, then other methods are available to ascertain whether polarisation effects are present. Firstly, the polarisation of the microscope illumination can be estimated. Here, we show that the intensities of the excitation lasers in the SoRa spinning disk confocal system used here vary sinusoidally according to the angle of a sheet polariser placed between the laser and a power meter. A cruder version of this is to simply visually observe whether the laser intensity varies when viewed through a polarising sheet that is rotated (of course while accounting for laser safety to avoid eye injury). Alternatively, imaging a sample in a circular dish and measuring orientation using AFT before and after manual rotation of the sample (as in Supplementary Figure S1) can help to disentangle real structural alignment from polarisation effects and indicate their relative strengths.

In addition to characterising polarised emission from fluorescently labelled collagen, we also present FouRD as a method for correcting inherent polarisation bias arising from an imaging system. We have demonstrated that FouRD can restore expected random orientations in AFT analysis of collagen matrices, and as FouRD corrects the original data it can be used successfully with other methods for orientation analysis (Supplementary Figure S3i, j). We would like to underline however that the performance of FouRD correction relies on performing imaging of the calibration sample in as similar conditions as possible to that of the data to be corrected; ideally the calibration sample would be made in the exact same way as the sample (except for the presence of any biological factors that could cause non-random alignment of the sample) and imaged during the same session as the test sample. This is because different reagent batches may polymerise differently, different types of sample dish may have different effects on polarisation (e.g. glass coverslip vs polymer coverslip), and there may be changes in microscope performance over time. For many biological structures and samples, as it is not technically possible to create a sample with random fibre orientations, we would instead suggest collecting a number of different fields of view covering a diverse range of structural organisation and averaging these to produce a calibration image. Although FouRD correction adds an extra experimental step, it is critical to avoid propagating systematic error through the analysis pipeline. Such errors can be particularly insidious when averaging over repeated experiments when increasing statistical power. As shown in Supplementary Figure S2, polarisation bias is particularly confounding for structures displaying mild alignment and is less disruptive to samples with strong alignment. Therefore, for proteins such as F-actin, where structures such as stress fibres display clear alignment that is readily perceptible to the human eye, FouRD correction is not essential as any orientational analysis will likely successfully detect this. However, the more subtle the change that is expected, the more critical it becomes that polarisation bias is compensated for.

## Conclusions

In this work we demonstrate that quantitative orientational analysis of fibrillar structures in fluorescence microscopy images can be disrupted by polarisation effects. These effects are seen for different fluorescently labelled fibrous structures, including collagen labelled with cyanine dyes via NHS ester linkage and actin labelled with Alexa Fluor 488 using phalloidin. This suggests that a range of fibre-forming biological structures may scaffold fluorescent labels in such a way that rotation is constrained or dipoles exhibit local alignment. While polarisation effects can be corrected using the FouRD method described here, these results in general should present a more cautionary tale for the application of quantitative image analysis to describe properties such as complex fibrillar orientation that are difficult to perceive by eye alone. Although researchers should not need in-depth knowledge of optics to use fluorescence microscopy, a basic understanding of potential artefacts that can arise is invaluable and can prevent biased results in the long term. This is particularly important in the age of automated image analysis, where algorithms may not be robust to uncommon but highly disruptive imaging artefacts. On a more positive note, should researchers encounter unexpected biases in automated analysis, this could indicate interesting properties of samples that could be exploited in developing future methods and protocols.

## Code and data availability

Code used for fibre simulation, Fourier Ring Anisotropy analysis and FouRD correction, along with data for testing, is available from https://github.com/CulleyLab/FouRD. This repository also contains a version of AFT code with small modifications were made to generate outputs such as the eccentricity maps.

## ACKNOWLEDGEMENTS

We would like to thank the staff at the KCL Nikon Imaging Centre, particularly James Levitt and Dylan Herzog, for their ongoing support. NA is funded by a Mechanics of Life Leverhulme Doctoral Scholarship, RM and SCu are funded by a Royal Society University Research Fellowship (URF\R1\211329), RM and SCo are funded by the Biotechnology and Biological Sciences Research Council (BB-SRC, grant no. BB/T011823/1), SM and BS are funded by the Wellcome Trust (grant no. 107859/Z/15/Z), the European Research Council (ERC) under the European Union’s Horizon 2020 research and innovation program (grant agreement no, 681808) and the BBSRC (grant no. BB/V006169/1).

## Notes

https://github.com/CulleyLab/FouRD

### Competing Interest Statement

The authors have declared no competing interest.

