## Supplementary Figures and Movie Captions for "Characterisation and correction of polarisation effects in fluorescently labelled fibres"

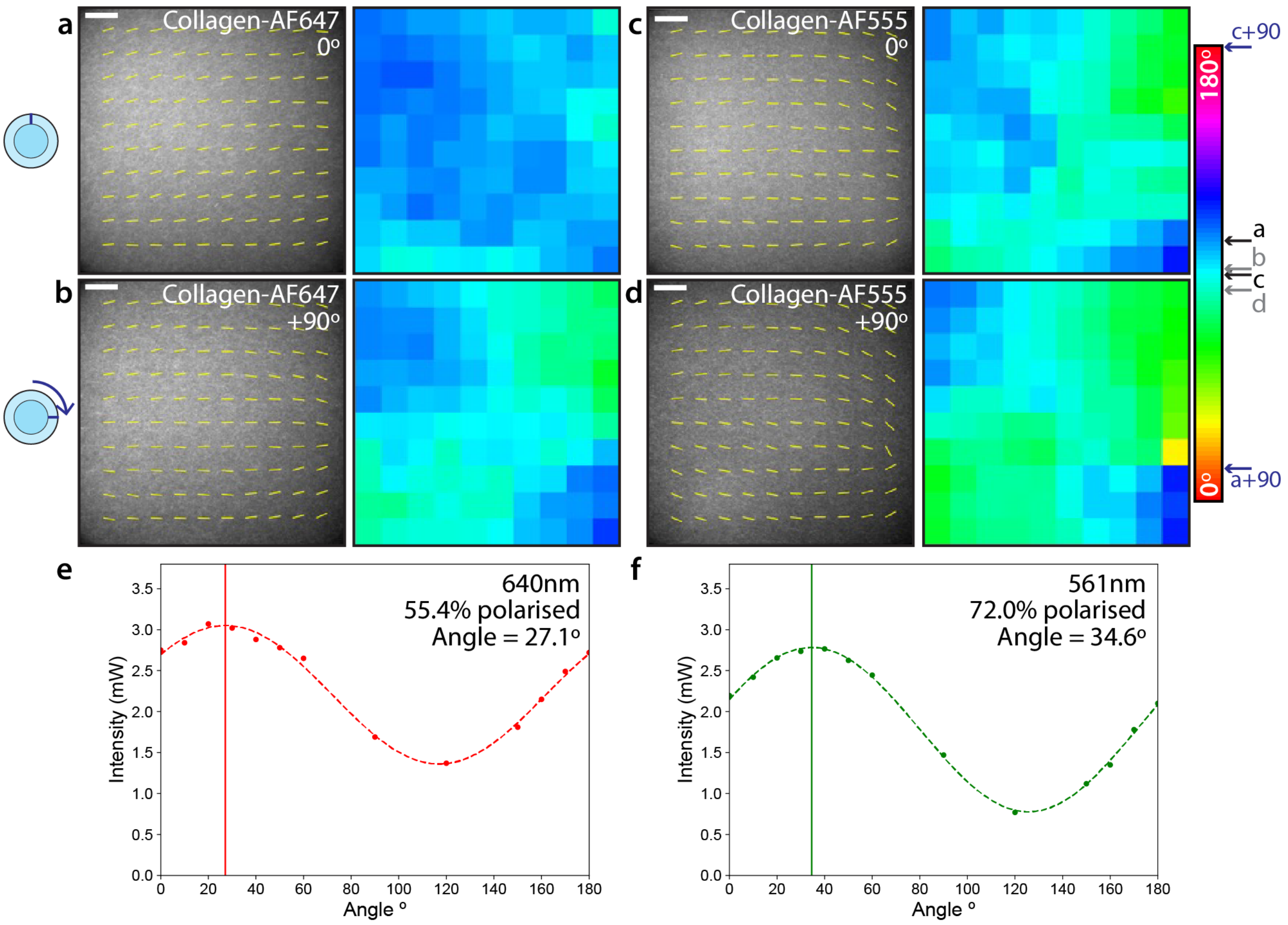


**Fig. S1. AFT analysis of physically rotated collagen gels shows persistent orientational bias.** Collagen-AF647 gel imaged on the SoRa spinning disk confocal microscope **a** before and **b** after physical ∼90º rotation of the sample dish, shown with the orientational vectors (yellow overlay) and angle colourmaps from AFT analysis. **c, d** Collagen-AF555 gel imaged on the SoRa spinning disk confocal microscope before (**c**) and after (**d**) physical ∼90º rotation of the sample dish shown with orientational vectors (yellow overlay) and angle colourmaps from AFT analysis. Vector maps from both gels display the same barrel-shaped orientation pattern before and after sample rotation. Mean (± standard deviation) alignments were **a** 100.1 ± 5.5º, **b** 90.8 ± 9.3º, **c** 88.5 ± 10.7º. **d** 82.5 ± 14.3º. Annotations on colour bar indicated mean measurements from samples before rotation (**a, c** - black) and after rotation(**b, d** - grey), as well as the expected mean angle if orientation was due to sample alignment (a+90, c+90 - dark blue). **e** Power measurements of the 640nm laser used for exciting collagen-AF647 taken at the back aperture when passed through a polarisation analyser at different orientations. Points indicate power measurements, dashed line indicates sine curve fitted to data. Solid line indicates location of peak angle. **f** As for **e**, but with the 561nm laser used for exciting collagen-AF555. Note that the angular frame of reference for the polarisation analyser is not the same as for the AFT analysis. AFT analysis was performed with 250 pixel window size, corresponding to 71.5 µm. Scale bars = 100 µm.

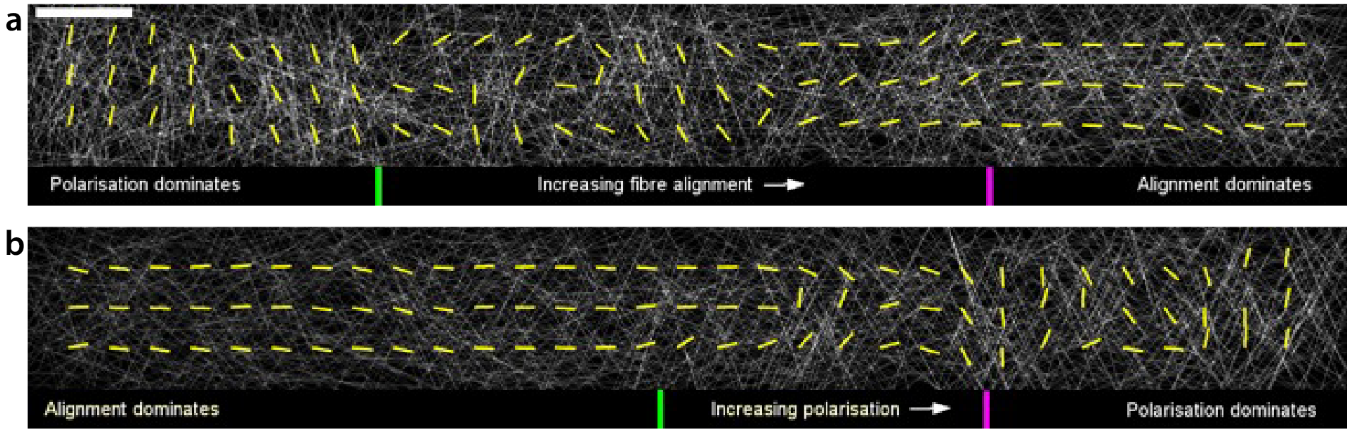


**Fig. S2. Simulated fibre networks with spatially varying degrees of alignment and polarisation indicate regimes where each effect dominates. a** Simulated fibres with alignment varying from left (totally random, 180º orientational range) to moderately horizontally constrained on the right (150º orientational range). Simulated excitation polarisation is constant at 50% vertically polarised, 50% unpolarised. Yellow overlay indicates AFT orientational vectors from AFT analysis. AFT analysis results to the left of the green marker are dominated by polarisation effects, whereas results to the right of the magenta marker are dominated by true alignment of fibres. **b** Simulated fibres with constant moderate horizontal alignment (within 150º orientational range) with simulated excitation polarisation varying from unpolarised on the left to 50% vertically polarised on the right. AFT analysis results to the left of the green marker are dominated by sample alignment, whereas to the right of the magenta marker they are dominated by polarisation effects. AFT block size was 250 pixels, corresponding to 25 µm.

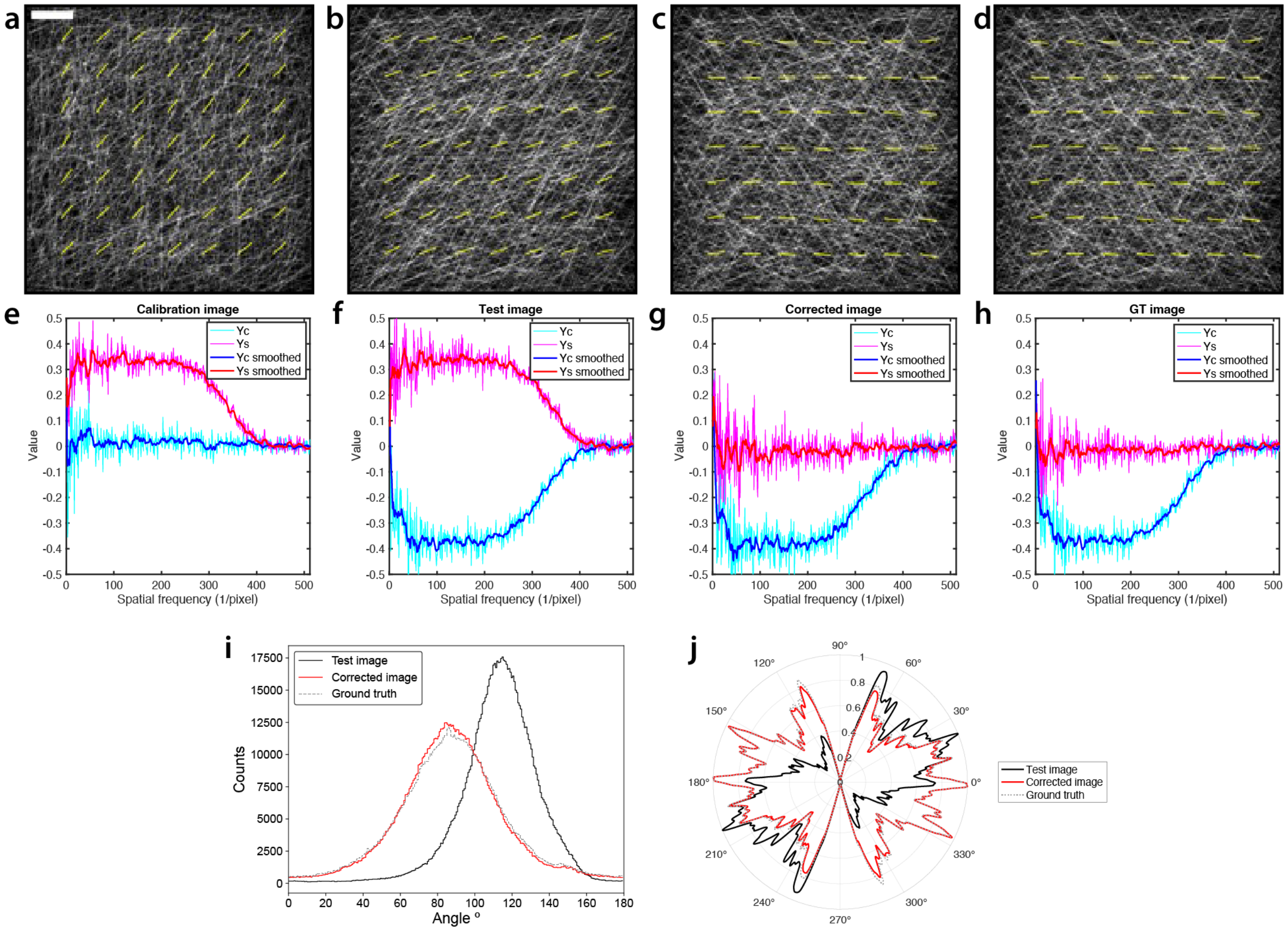


**Fig. S3. Demonstration of polarisation bias correction using FouRD on simulated data. a** To correct an image containing polarisation bias, we use a calibration sample containing randomly orientated fibres with simulated illumination with a polarisation operator of $1+\cos^2(\theta+45^\circ)$. This manifests in the yellow lines indicating direction vectors as calculated from AFT analysis. This also appears as a large $\sin 2\theta$ ($\gamma_s$) component in the anisotropy spectrum (**e**). This manifests in the yellow lines indicating direction vectors as calculated from AFT analysis. This also appears as a large $\sin 2\theta$ ($\gamma_s$) component in the anisotropy spectrum (**e**). **b** The target for polarisation correction is shown, which has an intrinsic alignment of fibres to horizontal ± 75º and the same polarisation operation as in **a**. The AFT vectors show a bias away from the expected 0º (horizontal) organisation. The anisotropy spectrum for **b** is shown in **f**, which shows significant $\cos 2\theta$ and $\sin 2\theta$ components. **c** The result of applying FouRD correction, based on the inverse of the anisotropy spectrum of the calibration image (**e**), to the test image in **b** shows a restoration of AFT vectors to 0º, and the corresponding anisotropy spectrum in **g** now shows a dominant $\cos 2\theta$ component. **d** The ground truth fibre distribution from the test image but without any polarised excitation. Strong agreement is seen between the FouRD-corrected image and spectrum (**c**, **g**) and the ground truth image and spectrum (**d**, **h**). **i** Histograms of orientation images generated by OrientationJ analysis of the images displayed in **b** (black line), **c** (red line) and **d** (dashed grey line). The angle histograms for the corrected and ground truth images are in excellent agreement. **j** CytoSpectre analysis of the images displayed in **b** (black line), **c** (red line) and **d** (dashed grey line). CytoSpectre reported the mean orientation angle in the test image as 27º, compared to 177º in both the corrected image and ground truth image. Again, the results of corrected and ground truth analysis agree very well. Scale bar = 10 µm. AFT block size was 250 pixels, corresponding to 25 µm.

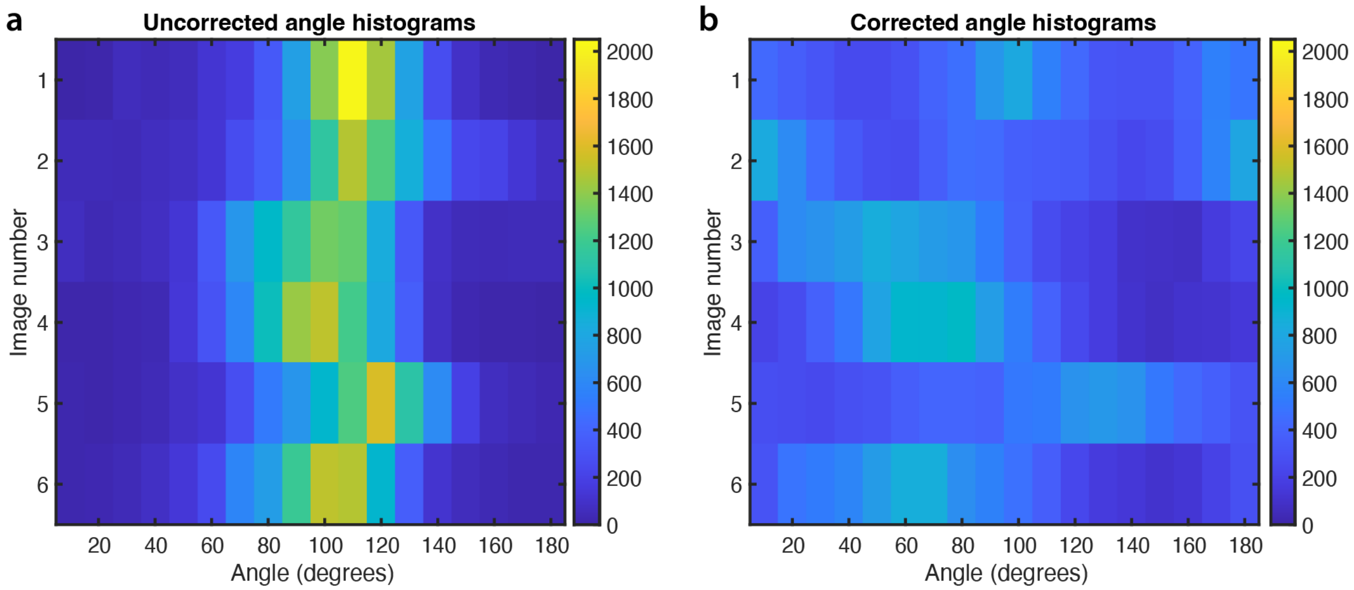


**Fig. S4. Aggregated results of AFT analysis applied to images before and after FouRD correction.** **a** AFT analysis was applied to 6 independent image z-stacks (76 slices each) of collagen-AF647 gels imaged on the SoRa microscope (each stack is a different row in the plot) and the measured angles summarised as histograms (colour indicates the number of pixels with orientation falling into the bins on the x axis). All histograms show a strong orientation peak in the 90-120º region, with little to no angles measured near 0º or 180º, due to polarisation. **b** FouRD correction was applied to the raw images analysed in **a** using the same calibration image in each case. AFT analysis was then performed and the angle histograms summarised and displayed in the same way as for **a**. Histograms of measured angles now have much less prominent peaks and are spread over a broader range of angles. The histogram peak locations are also now randomly distributed, demonstrating that the initial polarisation effect has been removed without adding any angle bias from the calibration. A perfect correction of a true random sample would yield a uniform histogram with a value of $\sim$530 in each bin. The SoRa data in Figure 6 corresponds to the first image of z-stack number 3 in this plot.

### Supplementary Movie 1:

**Graphical representation of the Fourier Ring Anisotropy calculation.** The process of determining the anisotropy is shown for a simulated fibre sample where each frame of the movie shows the calculation at an individual spatial frequency. The simulated fibres have no intrinsic orientational alignment but are subject to strongly polarised excitation (75%). The simulated horizontally and vertically polarised fluorescence images are shown in **a** and **b** respectively. The excitation polarisation in this case is sufficient for the orientational dependence of the images to be visible. The Fourier Anisotropy image $FA$ as calculated in Equation 3 is shown in **c** (quadrant swapped and log display scale). This image shows strong angular dependence for lower and mid spatial frequencies (towards the centre) but is rotationally isotropic at higher frequencies (towards the edges). This loss of anisotropy at higher frequencies is the combined effect of the finite point spread function (2.7 pixels FWHM) and noise. The yellow circle highlights all the indices in this image that correspond to the current spatial frequency, the 'Fourier ring'. To perform the calculation the sequence of values around this ring is used to form a 'profile' as a function of $\theta$, as shown by the black line in **d**. At lower frequencies the profile is dominated by a sinusoidal variation with a constant offset. This described the angular variation of the anisotropy image, which itself describes how the difference in brightness of the fibres between the two polarisation resolved images (**a** and **b**) varies with fibre orientation. The profile is then expanded into a Fourier series up to second order, the $\cos 2\theta$ (blue) and $\sin 2\theta$ (red) components of which are also plotted in **d**. At lower frequencies the $\cos 2\theta$ component is substantial, but at higher frequencies both components become small as the profile becomes flat and dominated by noise. The normalised magnitudes of the $\cos 2\theta$ and $\sin 2\theta$ are calculated (Equations 7a and 7b) to give anisotropy moments $\gamma_c$, $\gamma_s$ as a function of frequency. This gives the anisotropy spectrum shown in **e**, which indicates the strength and direction of any orientational bias to the emission.

### Supplementary Movie 2:

**Graphical representation of the FouRD correction process using a calibration image.** Each frame shows the correction performed using an additional spatial frequency, starting from the lowest frequency until the entire spectrum is included. **a** Real space image of the calibration image that has no intrinsic fibre alignment but does contain the effects of polarisation. **b** Absolute Fourier transform (log scale) of **a**. Taking the absolute removes real-space position information, leaving just orientation. The circle indicates all indices corresponding to the current spatial frequency. **c** Line profile (black) of the ring in **b** along with is $\cos 2\theta$ (blue) and $\sin 2\theta$ (red) components in a Fourier series expansion. **d** The normalised components $\gamma_c$, $\gamma_s$ as a function of frequency, the 'anisotropy spectrum', indicating orientational bias of the calibration image. **e** The inverse ring profile calculated from $\gamma_c$ and $\gamma_s$ indicating the opposing orientational bias needed at this frequency to remove the polarisation bias. **f** The inverse operator is constructed by adding the inverse profile (**e**) to the set of elements corresponding to the current spatial frequency (yellow circle, same set as in **b**). **g** The Fourier transform of the image to be corrected is multiplied (direct product) by the inverse operator **f** (log absolute shown in image). As all values in the inverse operator are real, it does not modify the position/structure (contained in the phase of the Fourier transform), just the relative brightness of certain orientations. **h** The corrected image is obtained from the inverse Fourier transform of **g**.
